# Comparative genomics analyses of lifestyle transitions at the origin of an invasive fungal pathogen in the genus *Cryphonectria*

**DOI:** 10.1101/2020.07.17.208942

**Authors:** Lea Stauber, Simone Prospero, Daniel Croll

## Abstract

Emerging fungal pathogens are a threat to forest and agroecosystems, as well as animal and human health. How pathogens evolve from non-pathogenic ancestors is still poorly understood making the prediction of future outbreaks challenging. Most pathogens have evolved lifestyle adaptations, which were enabled by specific changes in the gene content of the species. Hence, understanding transitions in the functions encoded by genomes gives valuable insight into the evolution of pathogenicity. Here, we studied lifestyle evolution in the genus *Cryphonectria*, including the prominent invasive pathogen *C. parasitica*, the causal agent of chestnut blight on *Castanea* species. We assembled and compared the genomes of pathogenic and putatively non-pathogenic *Cryphonectria* species, as well as sister group pathogens in the family Cryphonectriaceae (Diaporthales, Ascomycetes) to investigate the evolution of genome size and gene content. We found a striking loss of genes associated with carbohydrate metabolism (CAZymes) in *C. parasitica* compared to other Cryphonectriaceae. Despite substantial CAZyme gene loss, experimental data suggests that *C. parasitica* has retained wood colonization abilities shared with other *Cryphonectria* species. Putative effectors substantially varied in number, cysteine content and protein length among species. In contrast, secondary metabolite gene clusters show a high degree of conservation within the genus. Overall, our results underpin the recent lifestyle transition of *C. parasitica* towards a more pathogenic lifestyle. Our findings suggest that a CAZyme loss may have promoted pathogenicity of *C. parasitica* on chestnuts. Analyzing gene complements underlying key nutrition modes can facilitate the detection of species with the potential to emerge as pathogens.

## Introduction

Across the fungal kingdom, species have evolved the ability to persist either as symbionts, commensals or pathogens on a wide range of living insect, animal and plant hosts. This variety of fungal lifestyles requires complex adaptations encoded in the genome. Lifestyle-associated adaptations have been of particular interest as pathogen emergence is frequently associated with a significant gain in virulence of a formerly weak pathogen (Giraud et al., 2010). This has been shown for *Pyrenophora tritici-repentis*, a former saprophyte or weak pathogen on grass species including wheat, which became highly pathogenic on wheat through acquisition of the virulence gene *ToxA* from the wheat pathogen *Stagonospora nodorum* (Friesen et al., 2006). Moreover, pathogen emergence can be promoted through host jumps or geographic range expansions (Byrnes III et al., 2010) or complete host shifts (Giraud et al., 2010). Such host shifts can occur across kingdoms, as shown for insect pathogens from the genus *Metharhizium*, which likely evolved from plant endophytes or pathogens (Moonjely et al., 2016). Interestingly, phylogenomic analyses have shown that pathogens can emerge repeatedly within fungal clades such as *Dothideomycetes* or even at the genus level (*e.g. Aspergillus*) (Haridas et al., 2020; Rokas et al., 2020). Hence, many pathogenic fungi have non-pathogenic ancestors. This suggests that the emergence and evolution of pathogenic lifestyles is coupled with the acquisition of specific traits distinct from non-pathogenic relatives.

To be successful, pathogens must overcome physical and chemical barriers deployed by the host (Hardham, 2001). Plant pathogenic fungi have evolved specific lifestyles (*i.e.* biotrophy, hemi-biotrophy, necrotrophy) to exploit the host and each lifestyle requires distinct sets of genes (De Wit et al., 2012; O’Connell et al., 2012; Spanu et al., 2010; Stergiopoulos and de Wit, 2009). The gene repertoire of pathogens evolved through gene gains or losses, proliferation of transposable elements, as well as expansions or contractions of entire gene families, sometimes resulting in increased genome sizes, compared to related non-pathogenic species (Raffaele and Kamoun, 2012; Sánchez-Vallet et al., 2018). Gene families notably associated with fungal plant pathogenicity include enzymes for cell-wall degradation, small secreted proteins (*i.e.* effectors) and secondary metabolite gene clusters (Chooi and Solomon, 2014; De Jonge et al., 2011; Gaulin et al., 2018; Lelwala et al., 2019; Lyu et al., 2015; Macheleidt et al., 2016). Cell walls are an important physical barrier against pathogens but can be broken down and used as carbon sources by a variety of fungi. Carbohydrate active enzymes (CAZymes) specific for cellulose, hemi-cellulose or pectin degradation, are typically classified into the superfamilies of glycoside hydrolases (GHs), glycosyl transferases (GTs), polysaccharide lyases (PLs), carbohydrate esterases (CEs), as well as enzymes with auxiliary activities (AAs) and carbohydrate-binding modules (CBMs) (Kubicek et al., 2014). The types and number of CAZyme encoding genes varies among species and likely reflects adaptation to different nutritional niches (Glass et al., 2013). Most notably, necrotrophic pathogens tend to deploy cell-wall degrading enzymes to promote host damage and colonization (Rodriguez-Moreno et al., 2018). In contrast, biotrophic pathogens tend to have fewer enzymes involved in cell-wall degradation (Kubicek et al., 2014; Rodriguez-Moreno et al., 2018). Saprotrophic fungi feeding on decaying plant matter often show an overall reduced CAZyme complement compared to necrotrophic fungi (Hane et al., 2020), but specific expansions in CAZymes related to cellulose degradation (Ohm et al., 2012).

The emergence of pathogenic lifestyles has often required the ability to secrete effector proteins and secondary metabolites during contact with the host. Effectors are characterized as quickly evolving small, cysteine-rich secreted proteins, which are produced to manipulate plant host immune responses (Stukenbrock et al., 2011; Wang and Wang, 2017). Biotrophic and hemi-biotrophic pathogens secrete effector proteins to suppress host immunity and manipulate host cell physiology (Lo Presti et al., 2015). Necrotrophs deploy effectors also as host-specific toxins (Lo Presti et al., 2015; Tan et al., 2010). However, small secreted proteins resembling effectors are also expressed by saprophytic fungi and may be involved in degradative processes (Feldman et al., 2017). Virulence factors in pathogenic fungi can also include secondary metabolites, which are often low molecular weight compounds not essential for fungal growth. Polyketides, non-ribosomal peptides, terpenes and indole alkaloids are the main bioactive compounds acting as cytotoxins, antimicrobials or enzyme inhibitors (Keller et al., 2005). Genes underlying secondary metabolite biosynthesis pathways are often clustered in the genome (Nielsen et al., 2017). Secondary metabolites are produced by fungi of various lifestyles but may be more relevant virulence factors for necrotrophs, while biotrophs tend to lose the underlying genes (Spanu et al., 2010). Beyond pathogenicity-related functions, saprophytic or endophytic fungi produce secondary metabolites with important antimicrobial activity (Santos et al., 2017; Schalchli et al., 2011).

The family Cryphonectriaceae (Diaporthales, Ascomycetes) includes mainly bark-inhabiting species ranging from weak to severe pathogens (Gryzenhout et al., 2006; Jiang et al., 2020). The most aggressive pathogens include *Chrysoporthe* species affecting hosts in the order Myrthales (*e.g. Eucalyptus* spp.), as well as *Cryphonectria parasitica* (Murr.) Barr., the causal agent of chestnut blight on *Castanea* (Fagaceae) species (Gryzenhout et al., 2004; Rigling and Prospero, 2018). *C. parasitica* is native to East Asia (*i.e.* China, Korea, Japan) where it occurs as a weak pathogen on Chinese (*Ca. mollissima* Blume) and Japanese (*Ca. crenata* Siebold & Zucc.) chestnuts. However, *C. parasitica* was first described after its discovery in 1904 on American chestnut (*Ca. dentata* (Marsh.) Borkh.) in the United States (Rigling and Prospero, 2018). The rapid spread of the pathogen following its introduction resulted in the ecological extinction of *Ca. dentata* throughout its native distribution range in North America (Elliott and Swank, 2008). In Europe, chestnut blight was first observed in the 1930s and is nowadays present in all major chestnut growing areas (Rigling and Prospero, 2018). Following the colonization of Europe, *C. parasitica* has rapidly spread through most of southeastern Europe driven by the emergence of a highly successful lineage (Stauber et al. 2020). The invasion success likely stems from the establishment of a highly diverse European bridgehead population and a switch to asexual reproduction (Stauber et al. 2020). Besides host species in the genus *Castanea*, *C. parasitica* has been occasionally reported on oaks (*Quercus* spp.), maples (*Acer* spp.) and European hornbeam (*Carpinus betulus* L.) (Rigling and Prospero, 2018).

Both in the native and the invasive range, *C. parasitica* has closely related sister species, which are considered weak pathogens or saprophytes (Dennert et al., 2020). Among these, *C. japonica* Tak. Kobay. & Kaz. Itô (previously named *C. nitschkei*) was isolated from *Ca. crenata* in Japan (Liu et al., 2003; Myburg et al., 2004) and from oaks in China, on which it causes bark cankers (Jiang et al., 2019). The European species *C. naterciae* M.H. Bragança (syn. *C. decipiens*, Gryzenhout et al., 2009) was isolated from *Ca. sativa* and *Quercus* spp. in Portugal, Sardinia and Algeria (Bragança et al., 2011; Pinna et al., 2019; Smahi et al., 2018). Inoculation experiments showed that both *C. japonica* and *C. naterciae* are significantly less virulent on *Ca. sativa*, *Quercus robur* L. and *Fagus sylvatica* L. than *C. parasitica* (Dennert et al., 2020; Jiang et al., 2019;). Two other *Cryphonectria* species occurring in Europe are *C. radicalis* and *C. carpinicola*. The former is also present in North America and considered to be a saprophyte on dead wood of *Castanea* and *Quercus* species (Hoegger et al., 2002). Interestingly, the low prevalence may be the result of a displacement that occurred when the pathogenic sister species *C. parasitica* was first introduced to both continents (Hoegger et al., 2002). *C. carpinicola* is a recently described species isolated from declining European hornbeams in Austria, Georgia, Italy and Switzerland (Cornejo et al. pers. comm.). The complex lifestyles within the Cryphonectriaceae, including the emergence of new pathogens, raise important questions whether genetic factors facilitate pathogenic lifestyles.

In this study, we assembled and analyzed 104 genomes of the Cryphonectriaceae family including the major representatives *C. parasitica*, *C. radicalis*, *C. naterciae*, *C. japonica* and a recently detected European *Cryphonectria* species named *C. carpinicola* (personal communication). We analyzed orthology among the gene sets of the species and constructed a robust phylogenomic tree. We find that Cryphonectriaceae share similar trophic lifestyle traits. However, the chestnut pathogen *C. parasitica* has a substantially reduced complement in CAZymes. In contrast, the capacity to produce secondary metabolites is reduced among *Cryphonectria* species but is broadly conserved within the genus. Effector candidate proteins show genus and species specificity consistent with faster evolvability of the underlying genes.

## Materials and Methods

### Genome sequencing

We sequenced whole genomes of 90 *C. parasitica*, 3 *C. japonica*, 3 *C. radicalis*, 2 *C. naterciae* and 2 *C. carpinicola*. isolates covering the global distribution range (Supp. Table 1). All isolates were prepared for sequencing as described in Stauber et al. (2020). Sequencing was conducted using the Illumina HiSeq4000 and Illumina NovaSeq platforms (Illumina, San Diego, USA) at the Functional Genomics Center Zurich (FGCZ). By choosing the Illumina NovaSeq SP flowcell, NovaSeq reads were compatible with HiSeq4000 reads for downstream analysis. Several samples from Stauber et al. (2020) were included in this study. All sample accession numbers for the NCBI Short Read Archive are available in Supplementary Table S1. Outgroup genomes to the genus *Cryphonectria* were obtained for *Chrysoporthe cubensis*, *Chr. deuterocubensis* and *Chr. austroafricana* from NCBI BioProjects PRJNA279968, PRJNA265023 and PRJNA263707, respectively (Wingfield et al., 2015a, Wingfield et al., 2015b).

### Genome assembly and gene prediction

All 100 *Cryphonectria* spp. sequences were assembled with SPAdes v3.13.0 (Bankevich et al., 2012), using the --careful option and choosing the k-mers 21, 33, 45, 57 and 69 for the iterative assembly process. Genome sizes and assembly quality of *Cryphonectria* spp. *de novo* assemblies, as well as *Chrysoporthe* spp. draft genomes were assessed with QUAST v5.0.2 and BUSCO v3.0.2 (Mikheenko et al., 2018; Waterhouse et al., 2018). Gene models were predicted using BRAKER2 v2.1.4 (Hoff et al., 2019, 2016; Stanke et al., 2006, Stanke et al. 2008). Briefly, we set up gene annotation training using the existing *C. parasitica* v2 reference genome annotation (available from http://jgi.doe.gov/, (Crouch et al., 2020)) using the BRAKER2 options --alternatives-from-evidence=false, --fungus, --gff3 and --skip_fixing_broken_genes. For splice site hints, intron information was extracted from the reference genome annotation using the construct_introns function from the R package gread v0.99.3 (Srinivasan, 2019). After the training, genes were predicted in all assembled genomes using BRAKER2 adding coding sequence hints of the *C. parasitica* reference genome obtained using gffread v0.11.0 (Trapnell et al., 2012) and EMBOSS v6.6.0 tool transseq (Rice et al., 2011). We set the BRAKER 2 options --alternatives-from-evidence=false, --gff3, --useexisting, --prg=gth, --trainFromGth.

### Identification of orthologs and secondary metabolite gene clusters

To identify orthologs among all *Cryphonectria* and *Chrysoporthe* isolates we used OrthoFinder v2.3.7 (Emms and Kelly, 2019). We selected all single-copy ortholog groups and generated sequence alignments using MAFFT v7.429 (Katoh and Standley, 2013). Aligned sequences were used for phylogenetic tree building by generating 100 maximum-likelihood (ML) trees using the GTRCAT model with RAxML v8.2.12 (Stamatakis, 2014). The RAxML generated tree and bootstrap files were subsequently used to build a consensus tree with Astral v5.14.2 (Yin et al., 2019). The obtained consensus tree was visualized with FigTree v1.4.3 (Rambaut and Drummond, 2016). We used the antiSMASH fungal version v5.1.0 (Blin et al., 2019) to identify secondary metabolite gene clusters using one isolate per species; the reference genome EP155 for *C. parasitica*, IF-6 for *C. japonica*, M283 for *C. radicalis*, M3664 for *C. naterciae*, CS3 for *C*. *carpinicola* and the three NCBI *Chrysoporthe* draft genomes. We used a custom Python script to extract the biosynthetic core genes from the antiSMASH regions.js file. The number of core genes per species was then plotted in R with the packages tidyverse (Wickham, 2017), reshape2 (Wickham, 2007) and ggplot2 (Wickham, 2016). Additionally, we identified biosynthetic core genes per cluster of the EP155 genome and searched for orthologs in all species. Secondary metabolite core gene ortholog presence/absence in each cluster was plotted in R, using the packages reshape2, stringr (Wickham, 2018) and ggplot2.

### Classification of fungal lifestyles according to CAZyme content

We inferred trophic lifestyles of Cryphonectriaceae according to carbohydrate-active enzyme (CAZyme) gene content using the CAZyme-Assisted Training And Sorting of -trophy (CATAStrophy) prediction tool (Hane et al., 2020). CATAStrophy annotates CAZymes with HMMER 3.0 (Eddy, 2010) and dbCAN (Yin et al., 2012) and predicts trophic classes based on a multivariate analysis (Hane et al., 2020). To run CATAStrophy, we selected the same Cryphonectriaceae isolates as described above and added additional tree-associated fungi of different lifestyles. For non-pathogenic saprophytes associated with wood degradation, we selected the proteomes of *Fomitopsis rosea* (BioProject PRJNA518053), *Phanerochaete carnosa* (Suzuki et al., 2012) and *Phlebia centrifuga* (Mäkelä et al., 2018). Moreover, we included *Heterobasidion annosum* s.l. (Olson et al., 2012), associated with both saprotrophic and pathogenic lifestyles, and the bark pathogens *Neonectria ditissima* (Nectria canker on apple and pear trees) (Gómez-Cortecero et al., 2015), *Ophiostoma novo-ulmi* (Dutch elm disease) (Comeau et al., 2015; Forgetta et al., 2013) and *Valsa mali* (Valsa canker on apple trees) (Yin et al., 2015). For trophic lifestyle inferences, we used the catastrophy-pipeline (https://github.com/ccdmb/catastrophy-pipeline), choosing the options -profile conda and --dbcan_version 8. The CATAStrophy literature-derived nomenclature (*i.e.* classification into biotrophs, hemibiotrophs, nectrotrophs, saprotrophs and symbionts) was used for defining trophic lifestyles of species included in the CATAStrophy training set. Moreover, we selected principal components PC1 and PC2, which separate most training set species according to lifestyle for visualization (Hane et al., 2020).

### Analysis of carbohydrate-active enzymes genes (CAZymes) and inoculation experiments

For the identification of CAZyme genes we ran dbCAN v2.0.0 (Yin et al., 2012) on the same isolates as in the secondary metabolite analysis (*i.e.* one isolate per species). Only CAZymes which were identified by all three tools (HMMER, diamond, hotpep) were then selected for further analysis. CAZyme orthologs were extracted using Python and plots were generated in R.

To analyze the wood colonization capabilities of the different species, we set up an inoculation experiment. We selected 26 healthy, dormant chestnut logs (*Ca. sativa*; length: 50cm, diameter: 3.3-6.7cm), which were cut during winter from chestnut stands in Ticino, Switzerland, a week prior to the experiment. The logs were surface sterilized with 70% ethanol and sealed on both ends with paraffin to prevent desiccation. We selected 3 *C. parasitica* (XA19, CR03, EP155), 2 *C. japonica* (M9249, IF-6), 2 *C. naterciae* (M3664, M3656), 2 *C. radicalis* (M4733, M283), 2 *C*. *carpinicola* (M9290, CS3) and 1 *Chr. cubensis* (CBS115724) isolate for inoculation. For all isolates except CBS115724, full genome sequences were available for this study. Prior to inoculation, all isolates were freshly inoculated from glycerol stocks onto potato dextrose agar (PDA; 39 ml/L; BD Becton, Dickinson & Company; Franklin Lakes, USA) and incubated at 25°C in complete darkness for five days to induce mycelial growth. For inoculation of the first batch of chestnut logs (*n* = 13), we removed the bark on 5 equally distanced spots (diameter: 4mm) on each log, placed a mycelial plug (diameter: 4mm) into the wound and sealed it with tape. For the second batch of chestnut logs (*n* = 13), we directly placed five mycelial plugs onto the bark of each log with equal distance (*i.e.* no wound induction) and sealed the inoculation spots with tape. For each treatment batch with and without wound, we selected 5 replicates per isolate and 5 negative controls (mycelium-free agar plugs), resulting in a total of 130 completely randomized inoculation spots (*n* = 65 per treatment). All chestnut logs were randomly placed onto racks in plastic containers, separated by treatment, filled with 2L of demineralized water to avoid drying-out and sealed with plastic lids (Dennert et al., 2020). Incubation was at 20°C for both treatments. Logs with wounds were incubated for four weeks in complete darkness and longitudinal lesion size was assessed once a week. Logs without wounds were incubated for 12 weeks and lesion size was assessed at the end of the experiment.

### Prediction of effector genes

We performed effector gene prediction with the same isolates as in the secondary metabolite and CAZyme analysis, using a machine-learning approach. First, secreted proteins were predicted with SignalP v5.0b (Armenteros et al., 2019), choosing the options -org euk and -format short. Only proteins with a likelihood probability > 0.5 were selected for further analysis. Next, protein sequences with a predicted secretion signal were extracted with samtools v1.9 (Li et al., 2009) and used as input for effector prediction with EffectorP v2.0 (Sperschneider et al., 2018). Presence/absence variation analyses of predicted effector gene orthologs across species and plotting was performed in Python and R as described above. The cysteine content and protein length of predicted effector genes in all species were determined with EMBOSS pepstats v6.6.0.

## Results

### Genome assemblies for the *Cryphonectria* genus

We assembled draft genomes of 100 *Cryphonectria* spp. isolates of Asian, European and North American origin, in addition to the previously assembled genome of *C. parasitica* reference genome EP155. As a near outgroup to the genus *Cryphonectria,* we analyzed previously assembled draft genomes of 3 *Chrysoporthe* species from South Africa, Colombia and Indonesia. To assemble *Cryphonectria* genomes *de novo*, we used Illumina sequencing data at 9-53X coverage (Table 1). All *Cryphonectria* and *Chrysoporthe* genome assemblies showed >95% completeness for BUSCO genes (ascomycota_odb9 database) with the *C. parasitica* isolate M7832 having the lowest score at 95.9% (Table 1, Figure 1A). Based on the assembly size, we estimated that non-pathogenic species had smaller genomes ranging from 38.6Mb (*C. japonica)* to 41.9 Mb (*C. carpinicola*). Pathogenic species had slightly larger genomes ranging from 43.7 Mb (*C. parasitica* and *Chr. austroafricana*) to 45 Mb (*Chr. cubensis*; Table 1; Figure 1A). We found no apparent correlation between the estimated genome size and the completeness in BUSCO genes (Figure 1A-B). Similarly, we detected no correlation between the sequencing depth and the assembled genome size (Figure 1C). This shows that the short-read based assemblies are expected to reliably represent the gene content across species.

**Table 1:** Genome assembly statistics for *Cryphonectria* spp. and *Chrysoporthe* spp. The values correspond to mean values per species. The *C. parasitica* reference genome is not included in the summary of *C. parasitica* genomes.* = pathogenic species.

| Species | mean size [Mb] | mean N50 | mean complete BUSCO (%) | coverage (min-max) | N° predicted genes (min-max) | N° of isolates |
| --- | --- | --- | --- | --- | --- | --- |
| *C. parasitica* * | 43.7 | 125'044 | 98.40 | 9–53X | 11'321–12'195 | 91 |
| *C. japonica* | 38.6 | 364'390 | 98.43 | 17–48X | 10'680–10'729 | 3 |
| *C. radicalis* | 40.6 | 197'843 | 98.1 | 22–46X | 11'247–11'312 | 3 |
| *C. naterciae* | 39.4 | 120'703 | 98.05 | 19–27X | 11'041–11'050 | 2 |
| *C. carpinicola* | 41.9 | 58'390 | 97.35 | 14–26X | 11'159–11'187 | 2 |
| *Chr. austroafricana* * | 43.7 | 48'708 | 97.8 | NA | 13'125 | 1 |
| *Chr. cubensis* * | 45.0 | 345'702 | 96 | NA | 12'807 | 1 |
| *Chr. deuterocubensis* * | 43.9 | 83'661 | 96.2 | NA | 13'174 | 1 |

**Figure 1:**
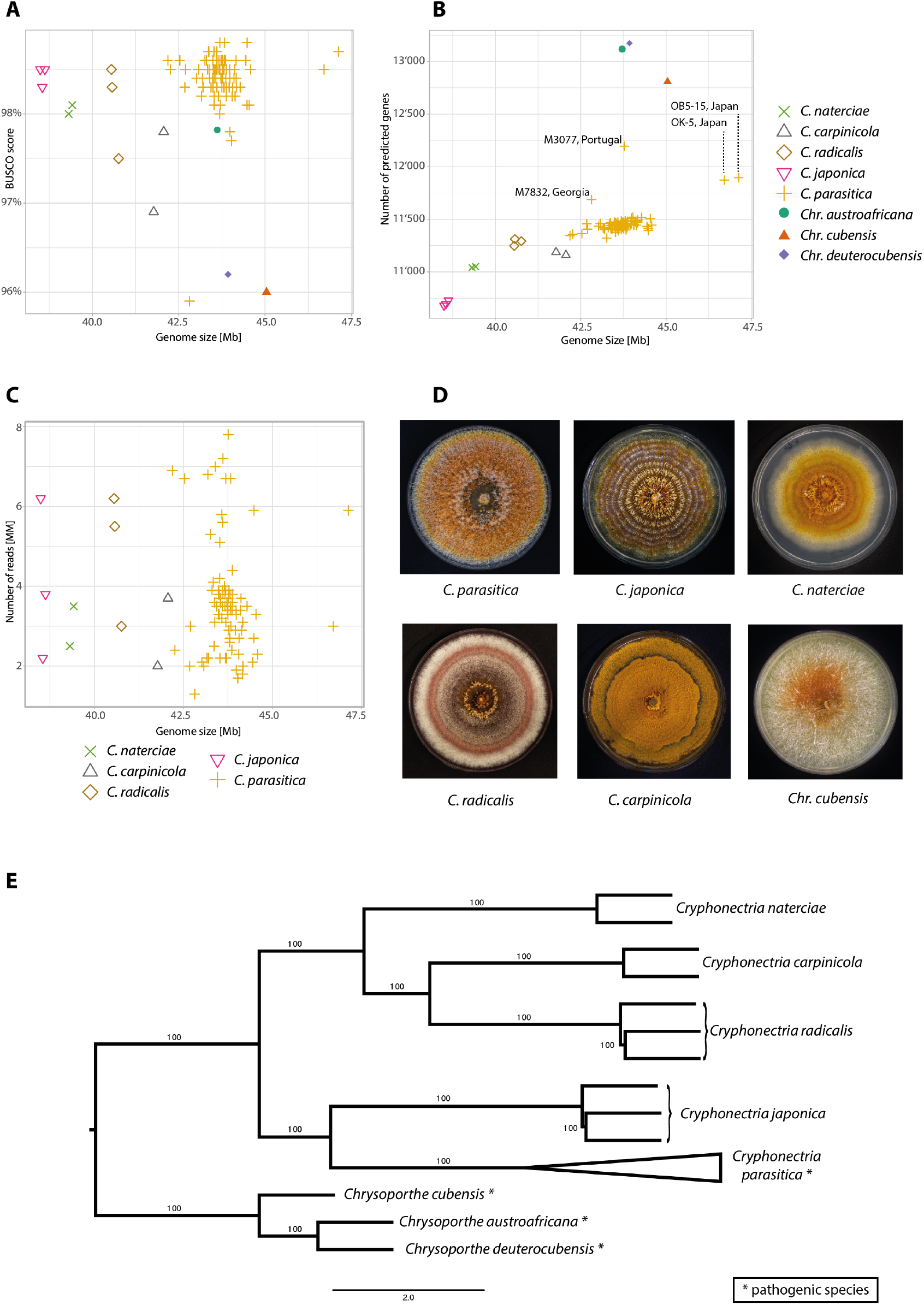
Assembly statistics and phylogenetic reconstruction. Estimated genome size in Mb correlated with A) assembly completeness assessed by BUSCO scores, B) number of predicted genes, and C) sequencing depth of assembled Cryphonectriaceae genomes. D) Culture morphology of the studied Cryphonectriaceae (cultures of *Chr. deuterocubensis* and *Chr. austroafricana* are not shown, as isolates were unavailable for documentation). E) Maximum-likelihood consensus tree based on 6770 single-copy ortholog genes showing the phylogenetic relationship of Cryphonectriaceae species.

### Gene annotation and phylogenetic reconstruction

We predicted between ~10’700–12’200 genes in genomes of *Cryphonectria* species compared to ~12’800 –13’170 genes in *Chrysoporthe* spp. (Table 1, Figure 1B). Overall, gene content among species was correlated with genome size except for *C. carpinicola* and *Chr. cubensis*, which have fewer predicted genes as expected from their genome size (Figure 1B). Among *C. parasitica* isolates, M3077 had a higher gene content compared with isolates of similar genome size (Figure 1B). Moreover, assembled genomes of *C. parasitica* isolates OB5-15 and OK-7 showed increased genome sizes while only having slightly higher gene content compared to other *C. parasitica* isolates (Figure 1B).

The gene ortholog analyses revealed 6770 single-copy orthologs among all species. We found 85 species-specific orthologs, of which 22 were specific for *C. parasitica*. Additionally, we found between 1–10 isolate-specific orthologs among the *C. parasitica* isolates TA51, M7832, DU5, OB5-15, OK-17 and M4030. Moreover, one ortholog was specific for *C. carpinicola*, while no species-specific orthologs were detected in all other *Cryphonectria* species. Within *Chrysoporthe*, we found 19 orthologs specific to *Chr. deuterocubensis*, as well as 12 and 5 orthologs specific to *Chr. cubensis* and *Chr. austroafricana*, respectively. To reconstruct the evolutionary history of *Cryphonectria* and *Chrysoporthe* species, we generated a consensus maximum-likelihood tree based on 6770 single-copy ortholog genes. We found 100% bootstrap branch support between species and a clear divergence at the genus level (Figure 1E). Furthermore, *Cryphonectria* species were grouping by geographic origin, with *C. naterciae*, *C. radicalis* and *C. carpinicola* being of European origin and *C. japonica* and *C. parasitica* being of Asian descent. Overall, our consensus tree is in accordance with phylogenetic studies on the genera *Cryphonectria* and *Chrysoporthe* (Cornejo et al., van der Merwe et al., 2010).

### Lifestyle prediction and capacity for carbohydrate metabolism across species

In order to degrade plant cell walls for nutrition or infection, fungi produce a variety of enzymes involved in carbohydrate metabolism (CAZymes) (Zhao et al., 2014). We analyzed the predicted proteome of Cryphonectriaceae species and other tree-associated fungi to assess trophic lifestyles according to CAZyme content. All *Cryphonectria* species were identified as hemi-biotrophs by CATAStrophy, while *Chrysoporthe* species were classified as necrotrophs. However, the PCA shows close proximity of analyzed *Cryphonectria* and *Chrysoporthe* species, clustering at the verge with other hemi-biotrophic and necrotrophic species (Figure 2A). Lifestyles of most fungi outside Cryphonectriaceae family matched with predicted lifestyles according to CAZyme content, except for *V. mali* (Figure 2A). The species is commonly defined as a necrotrophic pathogen (Li et al., 2015). However, in the CATAStrophy analysis *V. mali* clusters with other hemi-biotrophs and saprotrophs (Figure 2A). For *O. novo-ulmi*, phenotypic lifestyle classification remains uncertain (Sherif et al., 2016), but CATAStrophy classified this species as a symbiont or biotroph (Figure 2A), indicating a lower CAZyme content (Hane et al., 2020).

**Figure 2:**
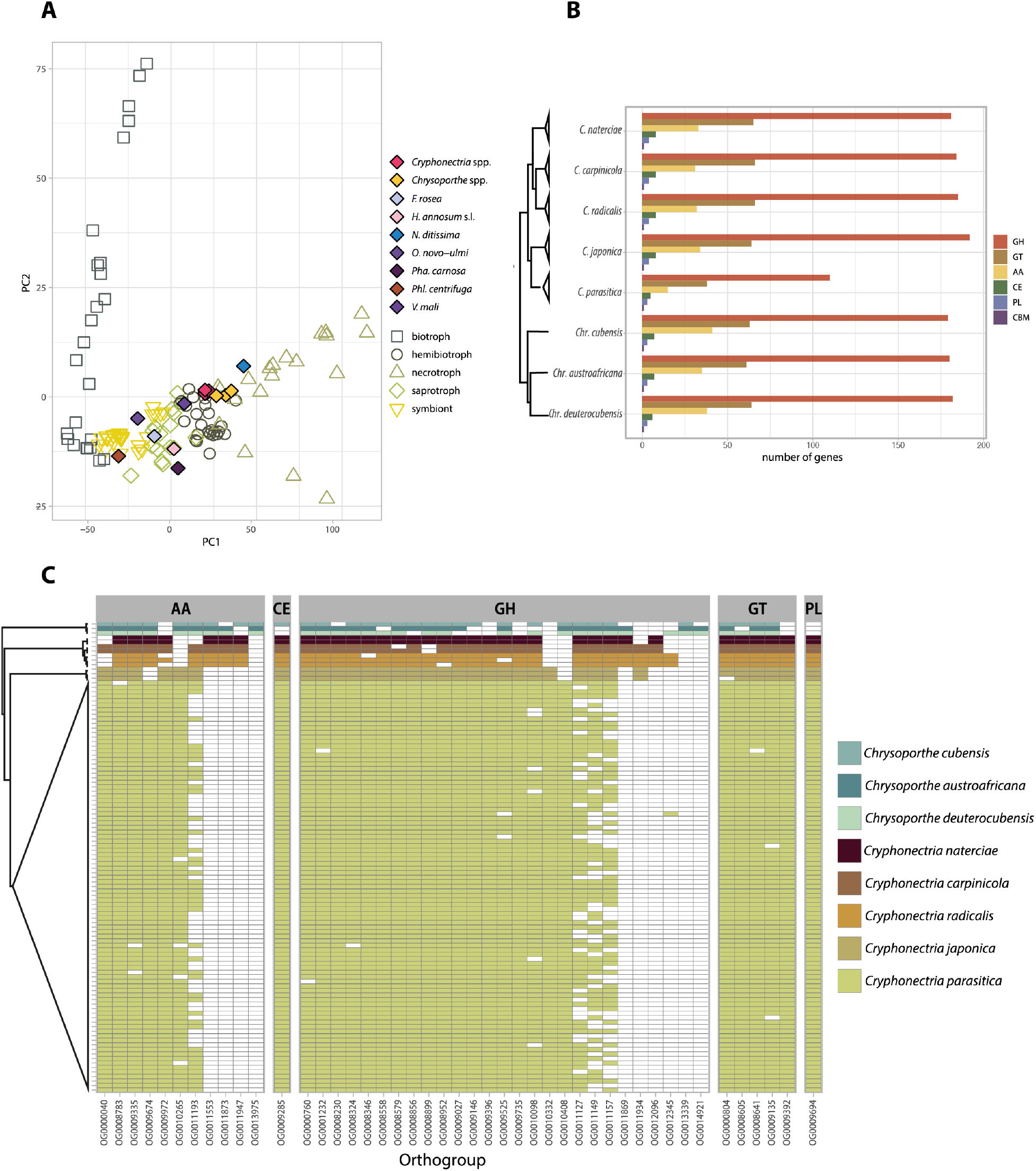
Carbohydrate Active EnZyme (CAZyme) content among Cryphonectriaceae. A) Principal component analysis (PCA) of fungal lifestyle predictions, as inferred by CATAStrophy. The plot incorporates 85 reference species of fungi with different lifestyles (*i.e.* biotroph, hemibiotroph, nectrotroph, saprotroph and symbiont) used as a training set by CATAStrophy and shows the CAZyme inferred phenotypic trophy of Cryphonectriaceae and other pathogenic and non-pathogenic tree-associated fungi. B) Number of detected CAZyme genes per species grouped according to CAZyme superfamily: glycoside hydrolase (GH), glycosyl transferase (GT), auxiliary activity (AA), carbohydrate esterase (CE), polysaccharide lyase activity (PL) and carbohydrate-binding modules (CBM). C) Ortholog presence/absence of CAZyme superfamilies for which at least one species is missing an ortholog (the CBM superfamily is not shown, as orthologs were found in all species).

We further assessed CAZyme gene content among Cryphonectriaceae and found a striking gene loss in the chestnut blight pathogen *C. parasitica* (Figure 2B). The gene loss particularly affected the group of glycoside hydrolases (GH), glycosyl transferases and enzymes with auxiliary activity (AA). Overall, all non-pathogenic *Cryphonectria*, as well as the pathogenic *Chrysoporthe* species encoded between 38.5– 42.7% more GH, 37.7–42.4% more GT and 51.6–63.4% more AA than *C. parasitica* (Figure 2B). We identified gene losses in *C. parasitica* across most CAZyme categories. GH5 associated with hemicellulose degradation showed a particularly remarkable reduction (Supp. Fig. 2). We found between 12–13 GH5 genes in saprophytic *Cryphonectria* and 11 GH5 genes in *Chrysoporthe* spp., while *C. parasitica* had only four GH5 genes. Moreover, slightly fewer GH28 genes involved in pectin degradation, were detected in *C. parasitica* (*n* = 11), compared to *Chrysoporthe* spp. (*n* = 12–14) and saprophytic *Cryphonectria* (*n* = 15–16). Analyzing CAZymes for which at least one species is missing an ortholog, *Cryphonectria* species share a relatively conserved set of ortholog CAZyme genes as expected from their short phylogenetic distance (Figure 2C). We found one PL orthogroup encoding for pectate lyase (OG0009694), only shared among *Cryphonectria* species. Moreover, we detected one GH orthogroup belonging to the sialidase superfamily (OG0010332), which is only present in Asian *Cryphonectria* species, as well as a single GH orthogroup (OG0012096: GH3) only present in European *Cryphonectria* species (Figure 2C). *C. parasitica* displayed a particularly high degree of intra-specific presence/absence variation for four auxiliary activity (AA) and GH enzymes, which are otherwise well conserved (OG0011193: GMC oxidoreductase, OG0011127: GH76, OG0011149: GH43, OG0011157: GH76; Figure 2C). The four orthogroups likely underwent recent gene losses in *C. parasitica.*

To assess the wood colonizing capabilities of different *Cryphonectria* species and a member of the genus *Chrysoporthe* (*Chr. cubensis*), we conducted an inoculation experiment on dormant chestnut stems. We performed the experiment with and without prior removal of the bark. None of the species were able to colonize dormant chestnut logs without artificial wound induction. After two weeks of incubation, *C. japonica* showed signs of mycelial growth on the bark at max. 1 cm beyond the inoculation point. No bark penetration was detected. For inoculations with bark removal, *C. parasitica* expectedly showed the fastest and most extensive lesion growth. Other *Cryphonectria* species, with the exception of one *C. radicalis* isolate (M4733), developed only minimal lesions (Supp. Fig. 1). We found intraspecific variance in lesion growth, possibly attributed to varying isolate vigor (*e.g. C. radicalis* isolate M283 was isolated in 1953) or varying substrate conditions (*e.g.* state of dormancy, stem thickness) (Supp. Fig. 1). The eucalyptus pathogen *Chr. cubensis* showed growth on non-host chestnut (*Ca. sativa*) logs, however, lesion developed at a comparatively slow pace (Supp. Fig. 1). After four weeks of incubation, mycelial fans were only found in lesions caused by *C. parasitica*.

### Variation in secondary metabolite production potential among species

Secondary metabolites (SM) can play important roles in pathogenicity and the interaction with microbes (Brakhage, 2013; Scharf et al., 2014). We investigated variation in biosynthetic core genes as an indicator for metabolite production potential among species. Loss of a biosynthetic core gene from a cluster invariably leads to loss of cluster function. Overall, biosynthetic core gene counts were only variable between genera. *Cryphonectria* species had comparatively fewer biosynthetic core genes than *Chrysoporthe* species (Figure 3A). Among the detected biosynthetic core genes, both genera shared similar proportions of different gene cluster classes with type 1 PKS (T1PKS) being the most abundant gene cluster class (Figure 3A). The class of beta-lactone production clusters, which can produce potent anti-bacterial and anti-fungal compounds (Robinson et al., 2019) were exclusively found in *Chrysoporthe* species (Figure 3A). The presence/absence analyses of biosynthetic core genes per gene clusters (*n* = 47) revealed 28 clusters conserved among all analyzed Cryphonectriaceae (Figure 3B). Additionally, core genes in five clusters were conserved in *Cryphonectria*. The same clusters showed partial or complete absence in *Chrysoporthe*. The largest cluster was found on scaffold 4, containing four T1PKS biosynthetic core genes. All four core orthologs of the cluster were retained in *C. parasitica.* Other species lost between one (*C. japonica*) and all four core genes (*Chr. cubensis*) (Figure 3B). Overall, core genes were highly conserved among *C. parasitica*, except for three T1PKS, NRPS-like and NRPS-T1PKS clusters on scaffolds 4, 6 and 11 (OG0010469, OG0005403 and OG0009181) (Figure 3B). The clusters showed gene losses in *C. parasitica* isolates from China (TA51), Georgia (M7776, M7832), Japan (WB-3) and USA (MD-1). Generally, we only identified weak homology with secondary metabolite clusters in other species. A notable exception includes two gene clusters potentially underlying emodin production (Supp. Figure 3).

**Figure 3:**
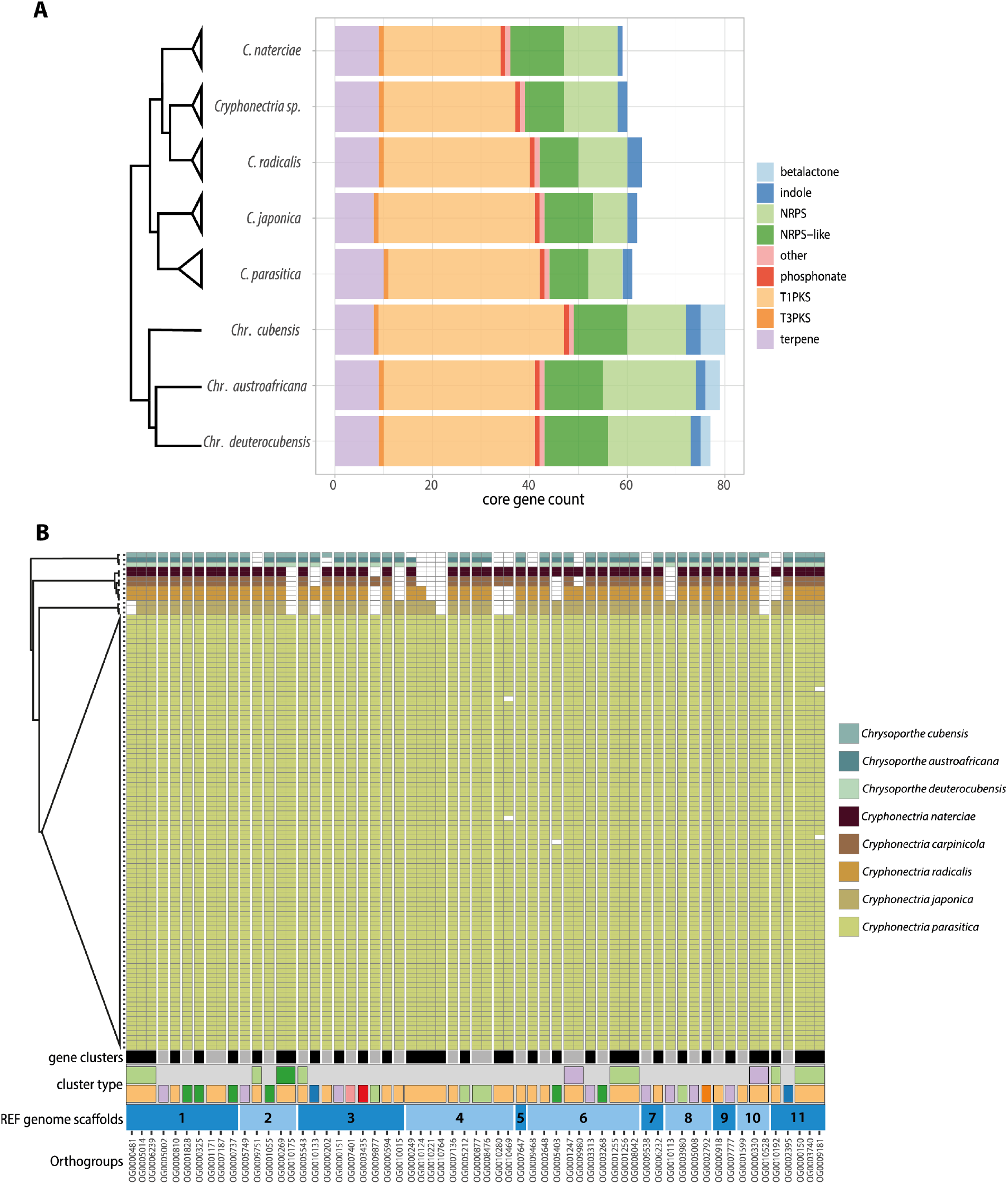
Secondary metabolite core gene content among Cryphonectriaceae. A) Count of detected biosynthetic core genes categories across species as identified by antiSMASH. B) Presence/absence of biosynthetic core gene orthologs among species. The plot shows the number of biosynthetic core genes within a gene cluster, the cluster type (color codes are as in A) and the location of clusters according to *C. parasitica* reference genome scaffolds.

### Predicted effector genes among Cryphonectriaceae, effector orthologs and cysteine content

Effectors are mostly secreted, cysteine-rich proteins, which play a major role in fungal virulence to overcome host immune defenses (Stergiopoulos and de Wit, 2009). We predicted effector genes with a machine-learning approach and found that neither the number of putative secreted proteins nor the predicted effector content correlated with genome size (Figure 4A). Saprophytic *Cryphonectria* species encode slightly more putatively secreted proteins (*n* = 777–796) compared to pathogenic *Chrysoporthe* spp. (*n* = 751–772). Surprisingly, *C. parasitica* encodes markedly fewer secreted proteins (*n* = 619) than all other species (Fig. 4A). However, despite the low amount of secreted proteins, *C. parasitica* had the highest ratio of predicted effectors among all species with 7.8% of all secreted proteins predicted to function as effectors (Figure 4A). Overall, the pathogenic versus saprophytic lifestyle did not correlate with predicted effector content. For example, we found that pathogenic *Chr. deuterocubensis* encoded the smallest number of predicted effectors of all analyzed species (Figure 4A). The cysteine content of predicted Cryphonectriaceae effectors ranged from 0–12.9% (Figure 4B). The predicted effectors among *Cryphonectria* contained 53–348 amino acids with one outlier of only 33 amino acids in *C. radicalis.* Predicted *Chrysoporthe* effectors contained 67–436 amino acids (Figure 4B). The divergence in candidate effector gene content among Cryphonectriaceae matches the divergence in cysteine content and protein length. Analysis of predicted effector ortholog presence/absence among Cryphonectriaceae revealed 41.5% (*n* = 59) conserved orthologs in all Cryphonectriaceae (Figure 4C). We found several orthologs unique to a single species (Figure 4C). Interestingly, the species-specific *C. parasitica* orthologs OG0010999, OG0010973 and OG0010938, showed presence-absence variation with orthologs missing in isolates from China and South Korea (LB86, M8510, S35; Figure 4C). For eight candidate effectors, we could not find a corresponding ortholog annotation with OrthoFinder (grey area in Figure 4C).

**Figure 4:**
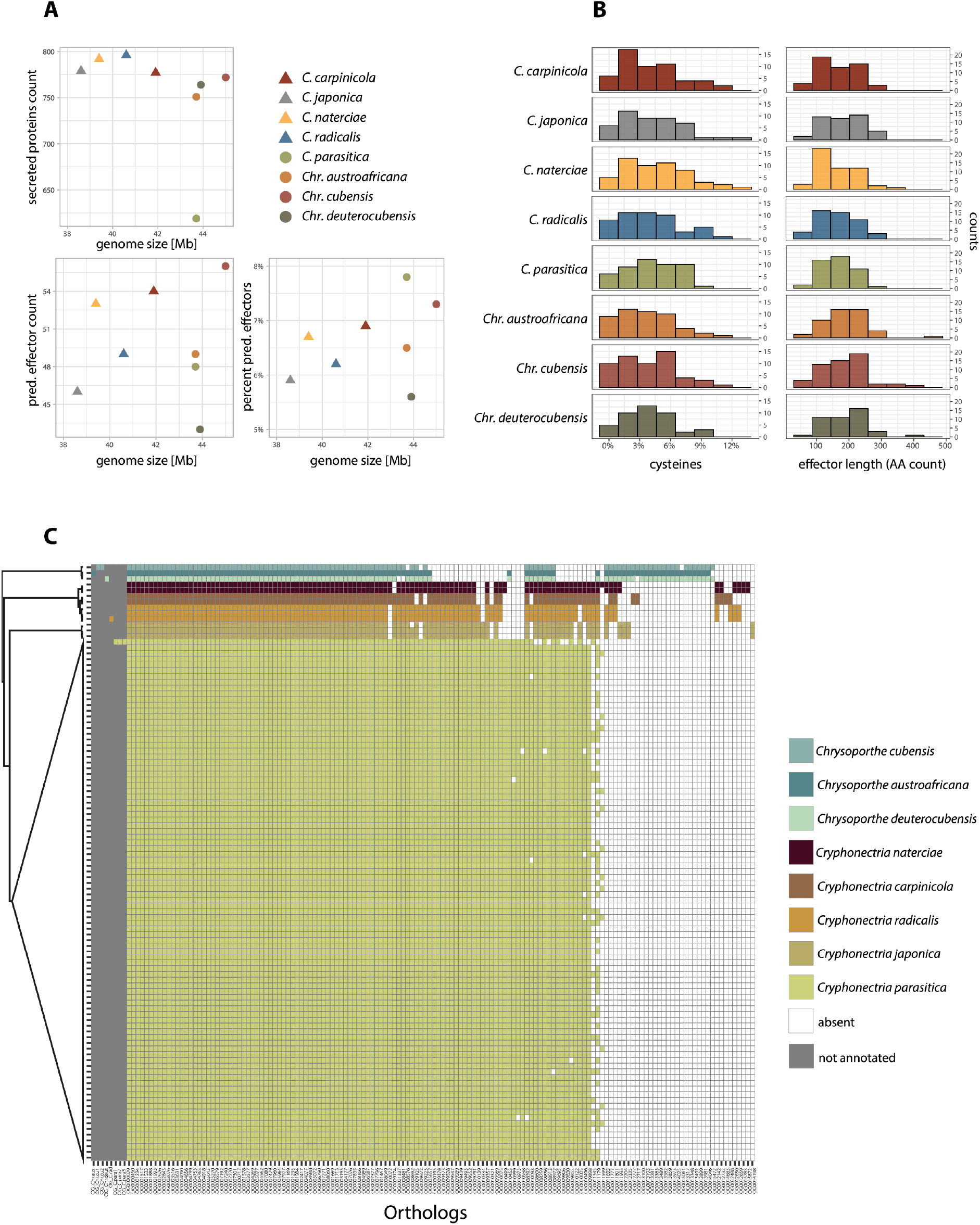
Predicted secretome and putative effectors among Cryphonectriaceae. A) Genome size correlations with secreted proteins and predicted effectors (identified by EffectorP). Saprophytic species are shown with triangles, pathogens with circles. B) Histograms showing the cysteine content (%) and the size of predicted effectors per species. C) Presence/absence of predicted effector orthologs among species. Areas in grey show orthologs for which we found no corresponding ortholog.

## Discussion

We assembled and analyzed genomes of eight bark-inhabiting Cryphonectriaceae species to retrace the evolution of genome size and gene content. Based on CAZyme content, all analyzed species are predicted to share a similar trophic lifestyle. In the genus *Cryphonectria*, we detected striking CAZyme gene loss in the invasive pathogen *C. parasitica*. This is consistent with a transition from a saprophytic to a predominantly pathogenic lifestyle within the genus. In spite of the substantial CAZyme gene loss, *C. parasitica* shares wood colonization strategies with the other *Cryphonectria* species and has retained the ability for early saprotrophic wood decay. In contrast, secondary metabolite gene clusters diverged at the genus level, but were largely conserved among *Cryphonectria* species. Putative effector content varied substantially among species with differences in cysteine content and protein length.

### Distinct CAZyme gene loss in a pathogenic species

Trophic lifestyle prediction according to CAZyme content classified all Cryphonectriaceae as having similar lifestyles, with CAZyme content being similar to other hemi-biotrophic or necrotrophic fungi. These results suggest that albeit substantial difference in pathogenicity (Dennert et al. 2020), Cryphonectriaceae species share trophic lifestyle traits. Thus, these results challenge previous classifications of *C. japonica*, *C. naterciae* and *C. radicalis* as predominantly saprotrophic species (Rigling and Prospero, 2018). Nonetheless, the distinct CAZyme loss in *C. parasitica* coincides with an increased pathogenicity towards non-native (*i.e.* non-Asian) *Castanea* species, which seems to be absent in other *Cryphonectria* species. Many CAZymes play a role in plant cell-wall degradation and can be important virulence factors in necrotrophic fungi. Reductions in CAZymes genes have been observed in biotrophic pathogens and are thought to be an adaptation to reduce the exposure of molecular patterns, which can trigger host defenses (Duplessis et al., 2014; Goulet and Saville, 2017). Moreover, CAZyme loss can occur during host shifts, such as from plant to animal or insect hosts (Zhang et al., 2018). In *C. parasitica,* the CAZyme loss may be an adaptation facilitating an increased pathogenic lifestyle. At the intraspecific level, fewer CAZymes are expressed during pathogenic growth compared to saprotrophic wood decay in the conifer pathogen *H. annosum s.l.,* which has plastic lifestyles (Olson et al., 2012). Similarly, *C. parasitica* may have undergone a transitory phase in the evolution of the predominant pathogenic lifestyle favoring reduced CAZyme expression and ultimately gene losses. Moreover, *H. annosum s.l.* produces more secondary metabolites including phytotoxins during the pathogenic lifestyle (Olson et al., 2012). These findings suggest that necrotrophic pathogens of trees have evolved different wood degradation strategies compared to saprotrophic relatives. The most significant gene loss in *C. parasitica* was found in the CAZyme subfamily GH5, which is underlying hemicellulose degradation. Consistent with this, GH5 expression is lower during pathogenic growth in *H. annosum s.l.* (Olson et al., 2012). In contrast, genomes of saprotrophic wood decayers such as *Pha. carnosa* have expanded GH5 repertoires (Suzuki et al., 2012). Despite extensive CAZyme loss in *C. parasitica*, our experimental data show that all *Cryphonectria* species, including *C. parasitica*, have retained similar wood colonization capabilities through bark wounds. Moreover, *C. parasitica* appears to have retained CAZymes suitable for early wood decay. This confirms field observations indicating that the fungus is able to survive a years on the bark of fresh dead chestnut wood (Prospero et al., 2006). In parallel to GH5, pectin degrading enzymes of GH28 are also slightly reduced in *C. parasitica.* However, polygalacturonases belonging to the GH28 family are suggested to contribute to virulence in *C. parasitica* (Lovat and Donnelly, 2019). Similarly, in other necrotrophic pathogens GH28 are also associated with pathogenicity showing expansions in the GH28 family (Sprockett et al., 2011). Similar to GH5, *C. parasitica* may have lost GH28 enzymes triggering host defenses through molecular pattern recognition by the host (Mengiste, 2012).

### Potential virulence-associated traits in *C. parasitica*

In contrast to the evolution of CAZymes, secondary metabolite production capabilities are largely conserved within the genus *Cryphonectria*. *C. parasitica* produces virulence-associated compounds including oxalic acid, tannases, laccases and phytotoxins such as cryparin and diaporthin but the genetic basis is only partially resolved (Lovat and Donnelly, 2019). The diaporthin production pathway is encoded by a PKS gene cluster in *Aspergillus oryzae* (Chankhamjon et al., 2016). However, we identified no clearly orthologous cluster in *C. parasitica*. The conservation of gene clusters across *Cryphonectria* species suggests that secondary metabolites played no particular role in the evolution of pathogenicity by *C. parasitica*. However, many fungi can modulate metabolite production depending on environmental conditions (Shwab and Keller, 2008). Hence, even if all *Cryphonectria* species share a core set of gene clusters, lifestyle transitions may induce differential expression depending on biotic or abiotic conditions. In addition to secondary metabolites, small secreted proteins (*i.e.* effectors) can play key roles in the emergence of new pathogens. We identified a broad pool of putative effector orthologs among Cryphonectriaceae. The size of the effector gene pool did not correlate with genome size or lifestyle as seen in other clades of plant pathogens (Lo Presti et al., 2015; Lowe and Howlett, 2012). Combining analyses of positive selection, gene expression and targeted gene deletion assays of effector candidates in *C. parasitica* will be needed to elucidate the role of effectors in causing chestnut blight.

### Lifestyle and the role of hosts

Cryphonectriaceae species represent a useful model to retrace how lifestyle transitions towards pathogenicity impact the evolution of gene content. On its native Asian hosts (*Ca. crenata* and *Ca. molissima*), the chestnut blight fungus *C. parasitica* causes only mild symptoms, which has been attributed to host-pathogen co-evolution (Rigling and Prospero, 2018). In contrast, on the naïve American and European chestnut species (*Ca. dentata, Ca. sativa*) the pathogen causes lethal bark cankers (Anagnostakis, 1992). In the invasive range, *C. parasitica* might also be a weak pathogen on *Quercus* spp., *Acer* spp. or *Carpinus betulus* (Rigling and Prospero, 2018). This suggests that *C. parasitica* has the genetic repertoire of a broad host range pathogen and that chestnut species may be the least able to resist pathogen invasion. In diverse forest ecosystems, disease incidence is often negatively correlated with host species richness (the “dilution effect”, Haas et al., 2011; Nguyen et al., 2017; Prospero and Cleary, 2017). Hence, growth on largely resistant hosts may be a bet hedging strategy of *C. parasitica* to survive and spread in the absence of the primary host (Trapero-Casas and Kaiser, 2009). The weak pathogenicity of other *Cryphonectria* species may be facilitated by environmental conditions, such as abiotic stress on the host or disturbance of the host microbiome (Brader et al., 2017). Subsequently, these *Cryphonectria* species may be considered as latent pathogens similar to some endophytes (Sakalidis et al., 2011; Vaz et al., 2018; Zakaria et al., 2010). Latent pathogenicity has been observed in other Cryphonectriaceae. For example, Granados et al. (2020) found that the *Eucalyptus* pathogen *Chr. cubensis* is an endophyte on Columbian Melastomataceae trees. Moreover, the pathogen *Chr. austroafricana* occurs as an endophyte in its native range but is pathogenic on non-native *Eucalyptus* trees (Mausse-Sitoe et al., 2016). Hence, host jumps likely facilitated the switch from endophytic to pathogenic lifestyle in both species (Granados et al., 2020; Mausse-Sitoe et al., 2016). *C. parasitica* has recently emerged as a major pathogen on non-Asian chestnut species. To what degree the extensive CAZyme loss increased the pathogenic potential prior to the emergence as an invasive pathogen remains to be investigated. Comparative genomics combined with gene function analyses provide a powerful approach to study lifestyle evolution and changes in the underlying genome architecture.

## Acknowledgements

We are grateful to Thomas Badet for helpful suggestions on a previous version of this manuscript. We thank Eva Augustiny, Silvia Kobel, Aria Minder, Quirin Kupper and Hélène Blauenstein for laboratory assistance. We acknowledge the Genetic Diversity Centre (GDC), ETH Zurich, and the Functional Genomics Center Zurich (FGCZ) for technical support and facility access. We also thank Martin Wrann for helping with photographic documentation. Paolo Cortesi, Michael Milgroom, Kiril Sotirovski, Mihajlo Risteski, Marin Ježić, Sang Hyun Lee and Seçil Akilli kindly provided samples. Michael and Brenda Wingfield gave valuable insight on phylogenetic relationships among Cryphonectriaceae. We kindly thank Sabina Moser Tralamazza for insightful discussions on secondary metabolites and for sharing scripts. LS was supported by the Swiss National Science Foundation (grant 170188 to SP).

