## Supplementary Figure for "Comparative genomics analyses of lifestyle transitions at the origin of an invasive fungal pathogen in the genus *Cryphonectria*"

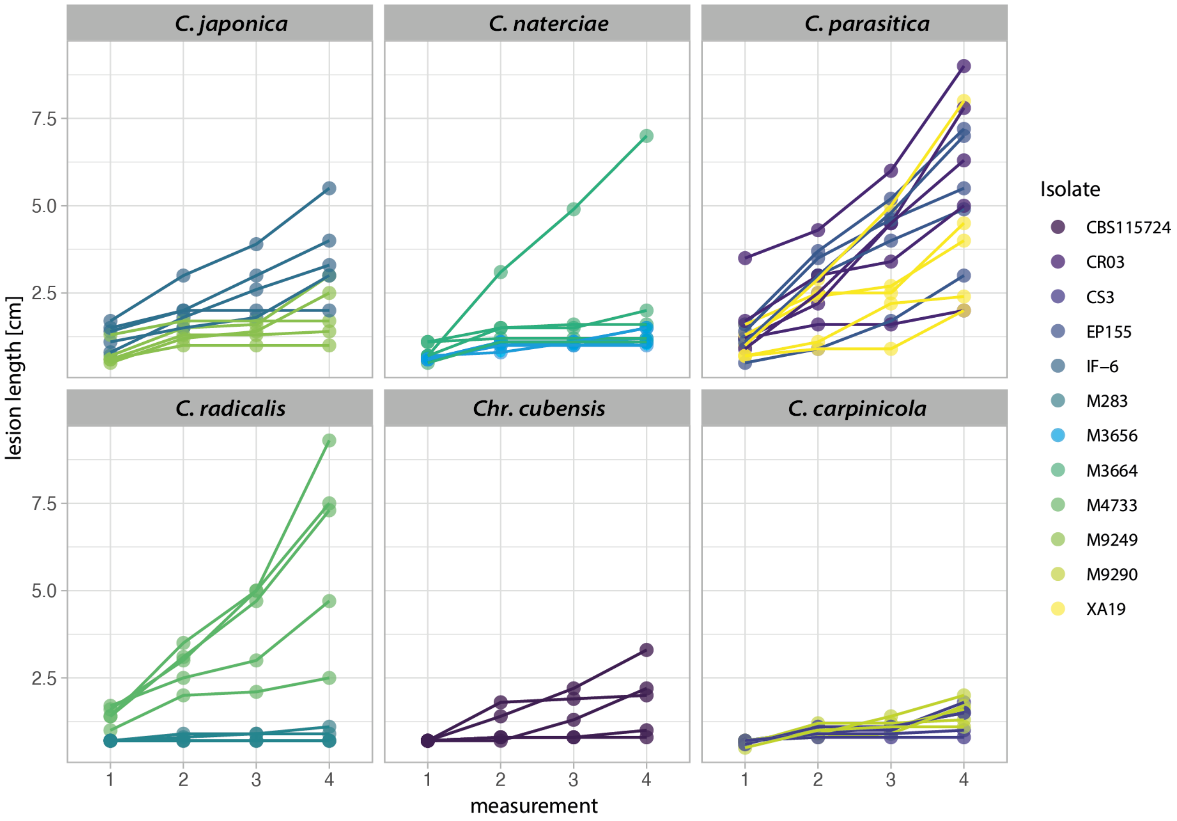

**Supplementary Figure 1:** Lesion length on dormant chestnut stems (*Castanea sativa*) with bark removal. Measurements were taken once per week during four weeks.

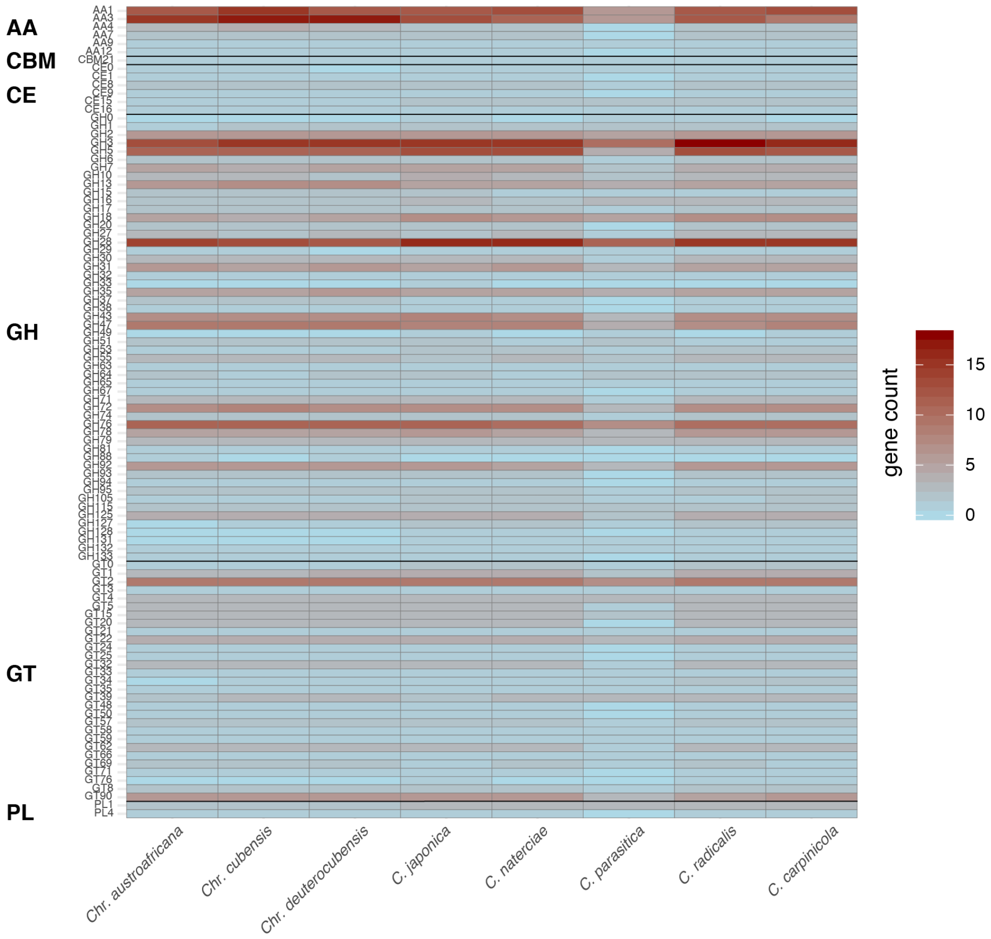


**Supplementary Figure 2:** Gene count of all identified CAZyme families among Cryphonectriaceae.

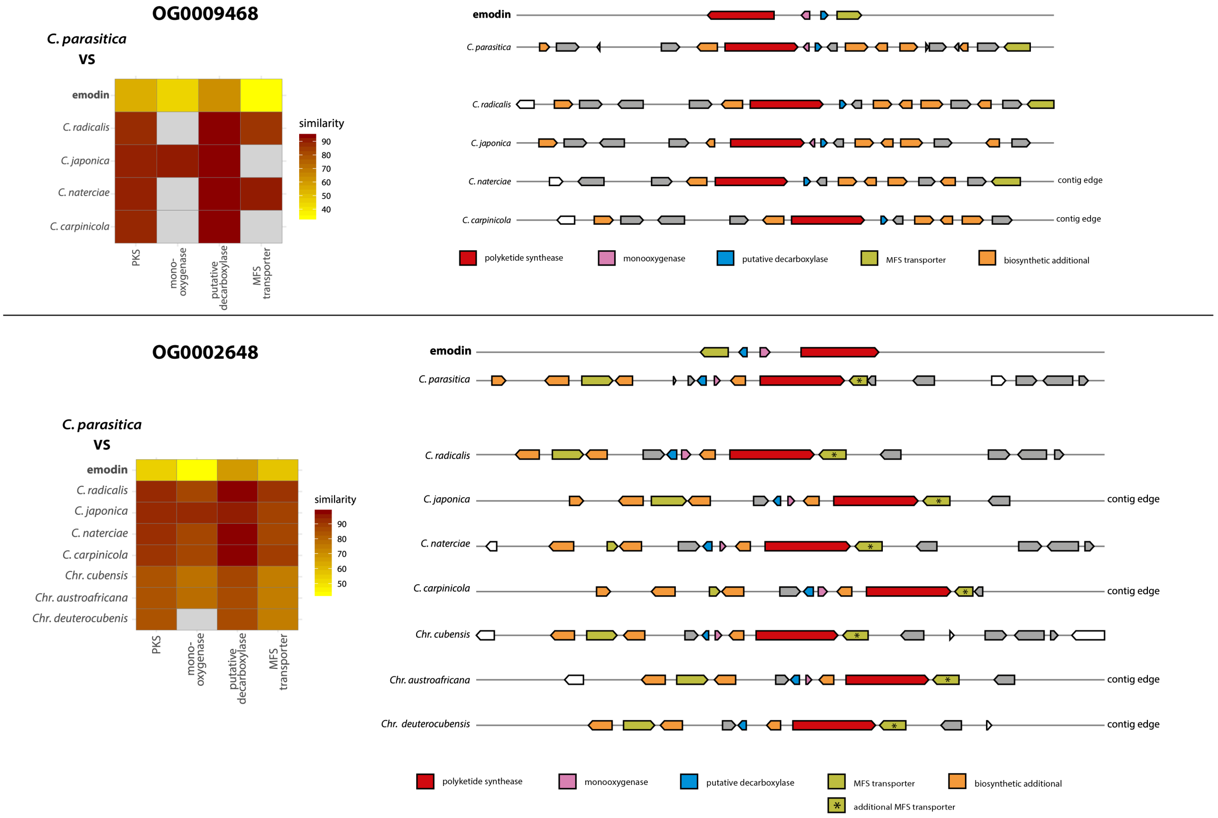


**Supplementary Figure 3:** Two PKS gene clusters (biosynthetic core gene orthologs OG0009468 and OG0002648) potentially underlying emodin production. Synteny plots of emodin and identified homologous gene clusters among species. Heatmaps show similarity (%) of genes in the *C. parasitica* reference genome cluster to emodin and orthologs in other Cryphonectriaceae (grey = gene is absent).
